# A mixed amplicon metabarcoding and sequencing approach for surveillance of drug resistance to levamisole and benzimidazole in *Haemonchus* spp

**DOI:** 10.1101/2023.05.14.540727

**Authors:** Emily Kate Francis, Alistair Antonopoulos, Mark Edward Westman, Janina McKay-Demeler, Roz Laing, Jan Šlapeta

**Author notes:** Author for correspondence: Jan Šlapeta.

## Abstract

Anthelmintic resistant parasitic nematodes present a significant threat to sustainable livestock production worldwide. The ability to detect the emergence of anthelmintic resistance at an early stage, and therefore determine which drugs remain most effective, is crucial for minimising production losses. Despite many years of research into the molecular basis of anthelmintic resistance, no molecular-based tools are commercially available for the diagnosis of resistance as it emerges in field settings. We described a mixed deep amplicon sequencing approach to determine the frequency of the levamisole (LEV) resistant single nucleotide polymorphism (SNP) within *arc*-8 exon 4 (S168T) in *Haemonchus* spp., coupled with benzimidazole (BZ) resistance SNPs within *β-tubulin* isotype-1 and ITS-2 nemabiome. This constitutes the first multi-drug and multi-species molecular diagnostic developed for helminths of veterinary importance. Of the ovine, bovine, caprine and camelid Australian field isolates we tested, S168T was detected in the majority of *Haemonchus* spp. populations from sheep and goats, but rarely at a frequency greater than 16%; an arbitrary threshold we set based on whole genome sequencing of LEV resistant *H. contortus* GWBII. Overall, BZ resistance was far more prevalent in *Haemonchus* spp. than LEV resistance, confirming that LEV is still an important anthelmintic class for small ruminants in New South Wales. The mixed amplicon metabarcoding approach described herein, paves the way towards the use of large scale sequencing as a surveillance technology in the field, the results of which can be translated into evidence-based recommendations for the livestock sector.

## 1. Introduction

Control of gastrointestinal nematodes (GINs) in livestock relies heavily on the availability of efficacious anthelmintics. Treatment failure, due to unregulated, frequent or incorrect application of broad-spectrum drenches targeting multiple parasites has resulted in anthelmintic resistance (AR), a global phenomenon now widespread in most major GINs of sheep, goats, and cattle (Charlier et al., 2022). Three important classes of anthelmintics are macrocyclic lactones (MLs) (ivermectin, abamectin, moxidectin), benzimidazoles (BZs) (albendazole, fenbendazole, oxfendazole), and imidazothiazoles such as levamisole (LEV). Development of resistance against LEV appears to be slower in comparison to MLs and BZs due to its lower frequency of use, which is why LEV remains an important ‘last resort’ anthelmintic in parasitic populations exhibiting multi-drug resistance (Tyrrell and LeJambre, 2010). Identification and surveillance of resistance markers at the molecular level is hence crucial for the sustainable utilisation of this drug in the field (Vercruysse et al., 2018).

Standard methods for detecting anthelmintic resistance in livestock are based on phenotype, whether by *in vivo* drug efficacy tests (i.e. faecal egg count reduction tests) or *in vitro* bioassays (i.e. egg hatch, larval motility or larval development assays) (Martin and Le Jambre, 1979; Coles et al., 2006; Demeler et al., 2012; Geurden et al., 2022). Whilst simple in theory, in reality these techniques are laborious, expensive and difficult to implement on a routine basis. Some lack the sensitivity required to detect early stages of resistance (i.e. when below 25% of the GIN population is resistant) (Martin et al., 1989; Levecke et al., 2011). In a post-genomic era, modern molecular diagnostic techniques that can detect known causal mutations of resistance, or genetically linked markers at low frequencies, for example measuring the frequency of SNPs in the *β-tubulin* isotype-1 gene of GINs to quantify resistance to benzimidazole, provide exciting opportunities for resistance surveillance and evidence-based management (Kwa et al., 1994; Silvestre and Cabaret, 2002; Kotze et al., 2012; Redman et al., 2015; Avramenko et al., 2019; Avramenko et al., 2020).

In *H. contortus*, LEV resistance has previously been associated with deletion mutations in the *acr-8* gene locus, yet recent data has implicated these as a poor markers of resistance due to their high frequency in LEV susceptible populations (Chagas et al., 2016; dos Santos et al., 2019; Baltrušis et al., 2021; Doyle et al., 2022). Using a genome-wide forward genetics approach with a genetic cross between susceptible MHco3(ISE) and multi-drug resistant MHco18(UGA) strains of *H. contortus,* a non-synonymous mutation, S168T, on *acr-8* exon 4 has since been identified as a key SNP in conferring decreasing sensitivity to LEV (Antonopoulos et al., 2022; Doyle et al., 2022; Baltrušis et al., 2023). LEV is a selective agonist of nematode nicotinic acetylcholine receptors (AChRs), of which *acr-8* is an important subunit (Boulin et al., 2011; Blanchard et al., 2018). Successful detection of the S168T variant in both laboratory and field populations of *H. contortus* was performed using allele-specific and droplet digital PCR, but requires optimisation for geographically divergent GIN isolates (Antonopoulos et al., 2022; Baltrušis et al., 2023).

In this paper, we developed a deep amplicon sequencing approach to determine the frequency of LEV resistant S168T in *Haemonchus* spp. in Australia, and coupled it with BZ resistance (*β-tubulin* isotype-1) and ITS-2 nemabiome deep amplicon sequencing for surveillance of parasitic nematode communities. Initially, we sequenced the whole genome of Australian LEV resistant and susceptible *H. contortus* and LEV susceptible *H. placei*. Based on the new genomes and those publicly available, we designed new primer sets targeting the *acr-8* exon 4 region encompassing the S168T SNP. The final assay was used to sequence a set of third stage larval (L3) samples from the field, either as individual amplicons, or mixed per field population. This approach shows the feasibility of concurrent rapid molecular detection of LEV and BZ susceptibility together with ITS-2 larval differentiation using metabarcoded deep amplicon sequencing.

## 2. Materials and methods

### 2.1. Parasite materials

#### 2.1.1. Resistant and susceptible Australian Haemonchus spp. L3 strains

Two Australian strains of *H. contortus* and one strain of *H. placei* were used in this study. 1) Kirby: *H. contortus* susceptible to all commercial anthelmintics; isolated from the University of New England Kirby Research Farm in 1986 (Albers and Burgess, 1988; Ruffell et al., 2018). 2) GWBII: Experimentally passaged *H. contortus* with demonstrated resistance to MLs, BZs and LEV (∼41% survival of adults post LEV treatment); a variant of multi-resistant GWBI isolated from Wallangra NSW in 2003 (Love et al., 2003; Kotze et al., 2018; Ruffell et al., 2018). 3) Grieve: *H. placei* susceptible to all commercial anthelmintics; isolated from a property in Enmore, NSW in 2017. All three strains were kindly provided by Invetus, Armidale NSW, in vials of ∼1000 L3 and had been cryopreserved prior to overnight transport to the University of Sydney on dry ice.

#### 2.1.2. L3 from ovine, bovine, caprine and camelid field isolates

Aliquots of >2,000 trichostrongylid L3s (N = 72) were provided by the Elizabeth Macarthur Agricultural Institute (EMAI, NSW Department of Primary Industries), Menangle NSW, Australia. L3s were cultured from ovine, bovine, caprine and camelid faecal samples submitted to EMAI from properties throughout NSW for routine gastrointestinal nematode diagnostic testing from May to September 2022. For each larval culture, the postcode, the date the sample was submitted for larval culture and the original EMAI morphological larval differentiation were available. Larval cultures were set up according to EMAI’s in-house procedures which are based on the Australian and New Zealand Standard Diagnostic Procedures (ANZSDP) (Hutchinson, 2009). Briefly, an equal slurry of fresh faeces and water was hand-mixed into vermiculite, until a slightly moist, but well-aerated mixture resulted. Cultures were incubated at 27°C for 7 days, to allow the development of L3, which were then harvested using a modified Baermann technique. The first 100 larvae were manually counted at 100× magnification to determine a larval percentage. L3 samples were transported at ambient temperature to the University of Sydney and stored at 4 °C until needed.

#### 2.1.3. DNA preparation

All parasite materials were centrifuged at 1,000 g for 2 min and total genomic DNA was isolated from the resulting pellets using the Monarch Genomic DNA Purification kit (New England Biolabs, Australia). For the ovine, bovine, caprine and camelid L3 field isolates, DNA was only extracted from 20 µL of the pellet, which equated to ∼2000 larvae. Isolations were conducted according to the manufacturer’s instructions for tissue lysis and ‘blank’ samples (ddH_2_O) were included in each batch to detect potential contamination during the isolation process. Each DNA isolate was eluted into a final volume of 100 µL genomic DNA elution buffer (10 mM Tris-HCl, pH = 9.0, 0.1 mM EDTA) and stored at −20 °C.

### 2.2. Sequencing of resistant and susceptible Australian *Haemonchus* spp. L3 strains

#### 2.2.1. Sanger sequencing

The *acr-8* exon 4 region encompassing LEV resistance marker S168T was first amplified from Australian *Haemonchus* spp. L3 strains (Kirby, GWBII and Grieve) using LEV1 and LEV2 primers which were designed based on the MHCO3ISE_4.0 *H. contortus* reference genome (PRJEB506) (Doyle et al., 2020). SYBR-chemistry real-time PCRs were performed using a Myra robotic liquid handling system (Bio Molecular Systems, Australia) at a final volume of 20 µL, containing SensiFAST™SYBR^®^ No-ROX mix (Meridian Bioscience, Australia), 2 µL of template DNA at a 1:1000 dilution, and primers at final concentrations of 400 nM. Reactions were run on a CFX96 Real-Time PCR Detection System with corresponding CFX Manager software (BioRad, Australia). The protocol involved an initial denaturation at 95 °C for 3 min, followed by 32 cycles of 95 °C for 5 s, 60 °C for 15 s, 72 °C for 15 s, and a final melt curve analysis. Each run included a no template control (ddH_2_O) to monitor for contamination. Amplicons were bidirectionally Sanger sequenced by Macrogen Ltd. (South Korea).

#### 2.2.2. Whole genome sequencing

DNA isolates from the Australian *Haemonchus* spp. L3 strains (Kirby, GWBII and Grieve) (40 µL: ∼50 ng/µL) were also sent to Novogene Co. Ltd. (Singapore) for indexing, library construction (350bp) and whole genome sequencing at an expected depth of 15 G raw data. Sequencing was performed using Illumina’s NovaSeq 6000 platform with 150bp paired-end reads (150PE). FastQ files containing the raw paired-end sequence reads were analysed using the free bioinformatics webserver, Galaxy Australia (https://galaxyproject.org) (Jalili et al., 2020). The inbuilt TrimGalore! function (Krueger et al., 2023) was performed for quality control and Illumina adapter trimming, followed by fast gapped-read alignment with Bowtie2 (Langmead and Salzberg, 2012) to map sequence reads to chromosome 5 of the MHCO3ISE_4.0 *H. contortus* reference genome (LS997566). Genome alignments were then visualised in IGV (Integrative Genomics Viewer) (Robinson et al., 2011) to inspect genetic variation around the *acr-8* exon 4 region to assist with primer design, and to confirm the LEV resistance status and proportion of S168T in the Kirby, GWBII and Grieve strains.

### 2.3. Design and validation of LEV *acr-8* primers for deep amplicon sequencing

#### 2.3.1. LEV acr-8 (S168T) primer design

We designed and tested multiple oligonucleotide pilot primer sets to amplify the LEV resistant S168T SNP within exon 4 of the *acr-8* genomic locus in *H. contortus* and *H. placei* (Table 1). To ensure performance on geographically divergent isolates, primers were designed against all available NCBI reference genomes for *H. contortus* (MHco3ISE, NZ_Hco_NP, McMaster) and *H. placei* (MHpl1), previous Sanger sequencing results of Australian adult *H.contortus* nematodes and L3 field isolates (see LabArchives for further details), in addition to the Sanger and whole genome sequencing of the Australian strains (Kirby, GWBII and Grieve). The forward primers were anchored in either intron 4 (LEV3, LEV 4, LEV5), similar to Antonopoulos et al. (2022), or exon 4 (LEV1, LEV6, LEV7, LEV8). Each primer differed due to between-strain polymorphisms and one primer at each binding site included multiple degenerate bases accounting for all of the observed variants. One reverse primer was designed (LEV2), which was conserved between all strains. All primers were manually designed in CLC Main Workbench (CLC Bio, Qiagen, Australia) and supplied by Macrogen Ltd. (South Korea) with Illumina adapters attached (Table 1).

#### 2.3.2. Validating LEV acr-8 (S168T) primers using Australian LEV susceptible and resistant Haemonchus spp. L3 isolates

To choose a final forward primer for future applications of the LEV *acr-8* metabarcoding technique, all primer sets were used to amplify exon 4 of *acr-8* including codon 168 from the Kirby, GWBII and Grieve isolates. SYBR-chemistry real-time PCRs were performed as described in *section 2.2.1*. The first 10 reactions were run on a 1% agarose gel stained with 0.1% GelRed (Biotium, USA) and visualised under a UV transilluminator to ensure amplicons were the required size. Amplicons were submitted for amplicon sequencing using Illumina next generation sequencing (NGS) at the Ramaciotti Centre for Genomics, University of New South Wales, Sydney, Australia. Here, stage-2 indexing PCRs, library quantification, normalisation, pooling, and denaturing were performed according to Illumina’s 16S Metagenomic Sequencing Library Preparation. Sequencing runs were undertaken on an Illumina MiSeq using MiSeq Reagent Kits v2 (500 cycles) for 250bp paired end (250PE) sequencing and de-multiplexed FastQ files were generated via BaseSpace (Illumina).

### 2.4. Combining LEV *acr-8*, BZ *β-tubulin* and ITS-2 nemabiome deep amplicon sequencing using L3 field isolates

LEV *acr-8* exon 4, BZ *β-tubulin isotype-1* and ITS-2 regions were amplified from ovine, bovine, caprine and camelid L3 field isolates (N=32) obtained throughout NSW. For the LEV *acr-8* assay, the degenerate exon 4 forward primer (LEV6) was selected, along with the reverse primer (LEV2). Both the ITS-2 and *β-tubulin* assays were performed using Illumina adapter primers previously described and validated (Avramenko et al., 2015; Avramenko et al., 2019). First-stage SYBR-chemistry real-time PCR’s were performed as individual assays (LEV, BZ, ITS-2) according to the protocol outlined in *section 2.2.1.,* with the exceptions that: 1) the final volume of each assay was 35 µL, 2) template DNA for the LEV and BZ assays was undiluted, whilst ITS-2 required a 1:1000 dilution, and 3) the PCR’s ran for 25-32 cycles (some amplicons required further cycling, likely due to differences in *Haemonchus* spp. DNA concentrations). To validate that the LEV, BZ, and ITS-2 amplicons for each field sample could be mixed prior to indexing and Illumina NGS without compromising sequencing results, of the 35 µL amplicon produced for each assay, 20 µL was first sequenced individually (standard protocol), and 10 µL was mixed via pipette with the other two assays prior to sequencing (mixed approach). An additional 40 L3 field isolates were screened for LEV S168T only using the LEV *acr-8* assay described above, performed at a final volume of 20 µL. Final mixed or individual un-indexed amplicons were then sent to the Ramaciotti Centre for Genomics for NGS within a 384 indexed amplicons using Illumina MiSeq.

### 2.5. Bioinformatic analysis

Sequence data and plotting of maps was performed in R (version 4.2.2). All LEV (*acr-8*), BZ (*β-tubulin*) and ITS-2 sequences (paired FastQ files) were analysed separately using the R package ‘dada2’ version 1.26.0 (Callahan et al., 2016) on Artemis HPC (Sydney Informatics Hub, The University of Sydney). In summary, primers were removed from all forward and reverse reads using ‘cutadapt’ version 4.0 (Martin, 2011) followed by ‘dada2’ pipeline including filtering, error estimation and denoising, before paired-end reads were merged to reconstruct the full amplicon sequence variant (ASV) and remove chimeras. LEV ASVs were assigned to the genus level using the available NCBI reference genomes for *H. contortus* (MHco3ISE, NZ_Hco_NP, McMaster) and *H. placei* (MHpl1). ITS-2 and BZ ASVs were assigned to the species level using in-house bespoke databases of local trichostrongylid nematode ITS-2 and *β-tubulin isotype-1* reference sequences curated from previously published data sets derived from the GINs of livestock in New South Wales, Australia (Francis et al., 2020; Francis and Šlapeta, 2022; Francis and Šlapeta, 2023). The databases were imported into ‘dada2’ using the ‘assignTaxonomy’ function, which assigns ASVs to the nearest match using the Bayesian classifier method, with a bootstrapping approach to assess the confidence assignment at each taxonomic level (Wang et al., 2007). Any ASVs not assigned to the species level using this approach were assigned using NCBI BLAST. All ASVs accounting for <0.1% of total reads across all samples were considered spurious and removed. Once AVSs were assigned to species or SNP variant those that accounted for <0.5% in a single sample were not considered further.

Shapefiles for Australia and postcodes were obtained from the Australian Bureau of Statistics (1270.0.55.001 - Australian Statistical Geography Standard (ASGS): Volume 1 - Main Structure and Greater Capital City Statistical Areas, July 2016, Australian Government). Shapefiles were processed and data from metabarcoding integrated using R packages ‘sf’ version 1.0-9 (Pebesma, 2018), ‘ggplot2’ version 3.4.1 (Wickham, 2016), and ‘dplyr’ version 1.1.0 (Wickham et al., 2023).

### 2.6. Statistical analysis and data accessibility

All ‘dada2’ pipeline output was analysed in Microsoft Excel (version 2207) and visualised in GraphPad Prism (version 9.5.1). Statistical analysis was performed in GraphPad Prism including two-way repeated measure (RM) ANOVA and specific details regarding the tests used are provided in the relevant sections of the results. Prior to statistical analysis, all datasets were tested for normal (Gaussian) distribution using the Shapiro-Wilk normality test. For all statistical tests, significance was set at *P* < 0.05. Raw FastQ sequence data was deposited at SRA NCBI BioProject: PRJNA962672. Local ITS-2 and *β-tubulin* isotype-1 trichostrongylid nematode databases and referenced supplementary data is available via LabArchives (https://dx.doi.org/10.25833/9h0d-0941).

## 3. Results

### 3.1. Sequencing of Australian LEV susceptible and resistant *Haemonchus* spp. confirms the presence of S168T and validates LEV NGS primer design

We used a combination of Sanger sequencing, WGS and deep amplicon NGS, to confirm the identity of the nucleotide sequence of amino acid residue 168 of *acr-8* exon 4 in Australian LEV susceptible and resistant *H. contortus* and susceptible *H. placei* L3 isolates, and to inform LEV NGS primer design (Fig. 1).

**Figure 1.**
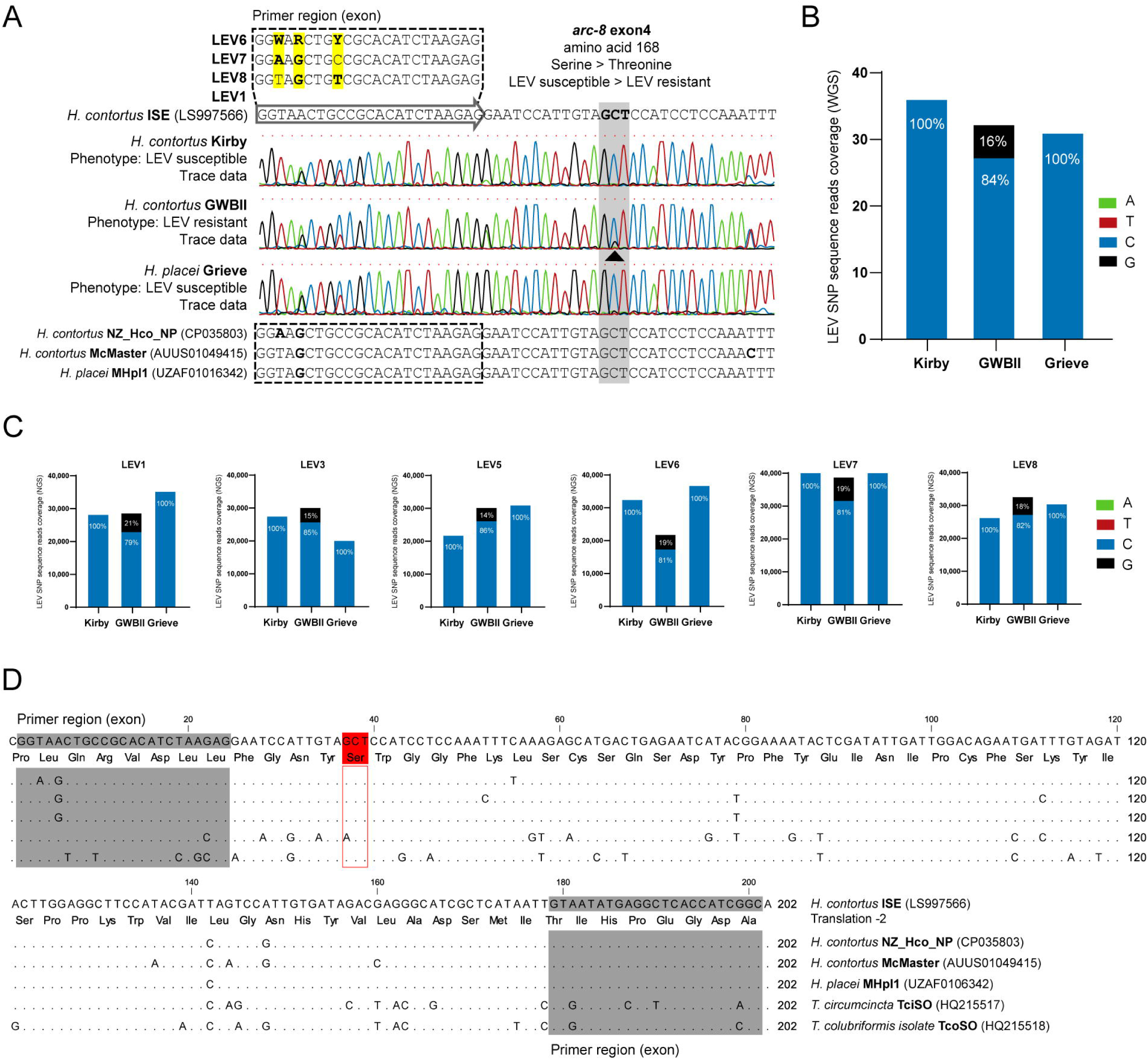
Confirmation and validation of *acr-8* exon 4 S168T in *Haemonchus* spp. metabarcoding assay. (A) Sanger sequencing of the arc-8 exon 4 of *Haemonchus contortus* from Kirby (levamisole susceptible) and GWBII (levamisole resistant) isolates, and *Haemonchus placei* Grieve (levamisole susceptible) isolate. The S168T (shaded) is detected only in GWBII as a secondary ∼1/5 peak (black arrowhead); note that S168T is coded on minus strand AGC>ACC. Primer region (dotted rectangle) shows rationale for degeneration of LEV6 primer due to polymorphisms in available genome data and/or the sequenced isolates. (B) Whole genome shotgun sequencing of the *Haemonchus* spp. DNA from Kirby, GWBII and Grieve isolates with detection and quantify of S168T variant conferring levamisole loss of efficacy. (C) Comparison of metabarcoded deep amplicon sequencing using six different combinations of primers to amplify and quantify S168T variant in *Haemonchus* spp. DNA from Kirby, GWBII and Grieve isolates. All amplification used LEV2 primer with LEV1, LEV3, LEV5-LEV8 (Table 1). (D) Multiple sequence alignment of the entire *acr-8* exon 4 for *Haemonchus* spp., *Teladorsagia circumcincta* and *Trichostrongylus colubriformis*. Translation (−2 coding frame) shows the conservation of amino acid sequence across all the sequences. Identical nucleotide sequences are represented as dots (.), primer regions are shaded and amino acid residue 168 (serine, AGC on minus strand) is highlighted.

Sanger sequencing of the phenotypically susceptible *H. contortus* Kirby and *H. placei* Grieve isolates produced unambiguous chromatographs with well-defined peak resolution at P168 (‘GCT’) analogous to reference *H. contortus* ISE (LS997566), confirming LEV susceptibility (Fig. 1A). The phenotypically resistant *H. contortus* GWBII produced a double peak at the second nucleotide of amino acid 168 with the minor peak being approximately 1/5^th^ the size and corresponding to the LEV resistant S168T genotype (‘GGT’). Chromatographs of all three isolates also confirmed variation in respect to reference *H. contortus* ISE (LS997566) within the exon 4 forward primer binding site, analogous to the geographically divergent variants observed between the other available reference genomes (CP035803, AUUS01049415, UZAF01016342). Accordingly, all variants and corresponding degenerate bases were accounted for when designing the seven pilot LEV NGS primer sets (Table 1).

WGS of the Kirby, GWBII and Grieve isolates yielded an average reading depth of 17G raw data (15.9 – 17.9) and 114,316,562 raw reads (106,159,252 – 119,310,774). Each isolate was successfully aligned to chromosome 5 of the *H. contortus* ISE reference genome (LS997566), revealing 100% LEV susceptibility at *acr-8* exon 4 P168 for *H. contortus* Kirby (100% G<u>C</u>T, 36: 18+, 18-) and *H. placei* Grieve (100% G<u>C</u>T, 31: 11+, 20-), and 16% LEV resistant S168T for *H. contortus* GWBII (84% G<u>C</u>T, 27: 15+, 12-; 16% G<u>G</u>T, 5: 1+, 4-) (Fig. 1B). The 16% G<u>G</u>T was directly proportional to the minor peak observed for resistant S168T on the Sanger chromatograph. WGS of each *Haemonchus* spp. isolate was also aligned to chromosome 1 of the *H. contortus* ISE reference genome (LS997562) to measure BZ resistance at *β-tubulin isotype-1* P167, P198 and P200, see LabArchives (https://dx.doi.org/10.25833/9h0d-0941).

Seven pilot LEV NGS primer sets with forward primers flanking either intron 4 or exon 4 encompassing *acr-8* S168T, were validated via next generation amplicon sequencing of the Kirby, GWBII and Grieve isolates (Fig. 1C; Table 1). All primers yielded a positive SYBR-chemistry real-time first-stage PCR result (C_t_ < 32) for both *H. contortus* and *H. placei* isolates (mean C_t_ = 25.8, SD 4.1) with the exception of LEV4 that was not considered further. Following Illumina MiSeq (250PE) NGS and ‘dada2’ analysis, the remaining six primers all produced >20,000 reads for each isolate (mean = 30,922, SD 6,743) which were processed into a total of 71 unique ASVs. SNP analysis showed all primers successfully detected LEV resistant S168T in *H. contortus* GWBII in similar proportions to the 16% revealed via WGS (LEV1: 21%, LEV3: 15%, LEV5: 14%, LEV6: 19%, LEV7: 19%, LEV8: 18%), whilst *H. contortus* Kirby and *H. placei* Grieve remained LEV susceptible (Fig. 1C). The *acr-8* exon 4 was highly conserved between the two species with no discernable nucleotide difference across the exon 4 sequence between *H. contortus* and *H. placei* isolates (Fig. 1D). Although all six primers were successful LEV NGS candidates, we decided to move forward with LEV 6; the degenerate exon 4 primer due to its universality.

In the highly LEV resistant GWBII isolate, the G<u>G</u>T mutation encoding S168T was present in 16% WGS reads aligned to the *acr-8* locus. This proportion was therefore used as an arbitrary threshold for LEV resistance for the remainder of our study.

### 3.2. Rapid molecular detection of LEV and BZ resistance, together with ITS-2 larval differentiation, using metabarcoded deep amplicon sequencing

To validate a rapid mixed molecular approach that simultaneously detects larval isolates to the species level and surveils LEV and BZ resistance, we obtained 32 field isolates of L3 of ovine, bovine, caprine and camelid origin. ITS-2, *acr-8* exon 4, and *β-tubulin isotype-1* regions were amplified and amplicons were first sequenced individually, and then mixed at a 1:1:1 ratio, using Illumina NGS (Fig. 2; Fig. 3A).

**Figure 2.**
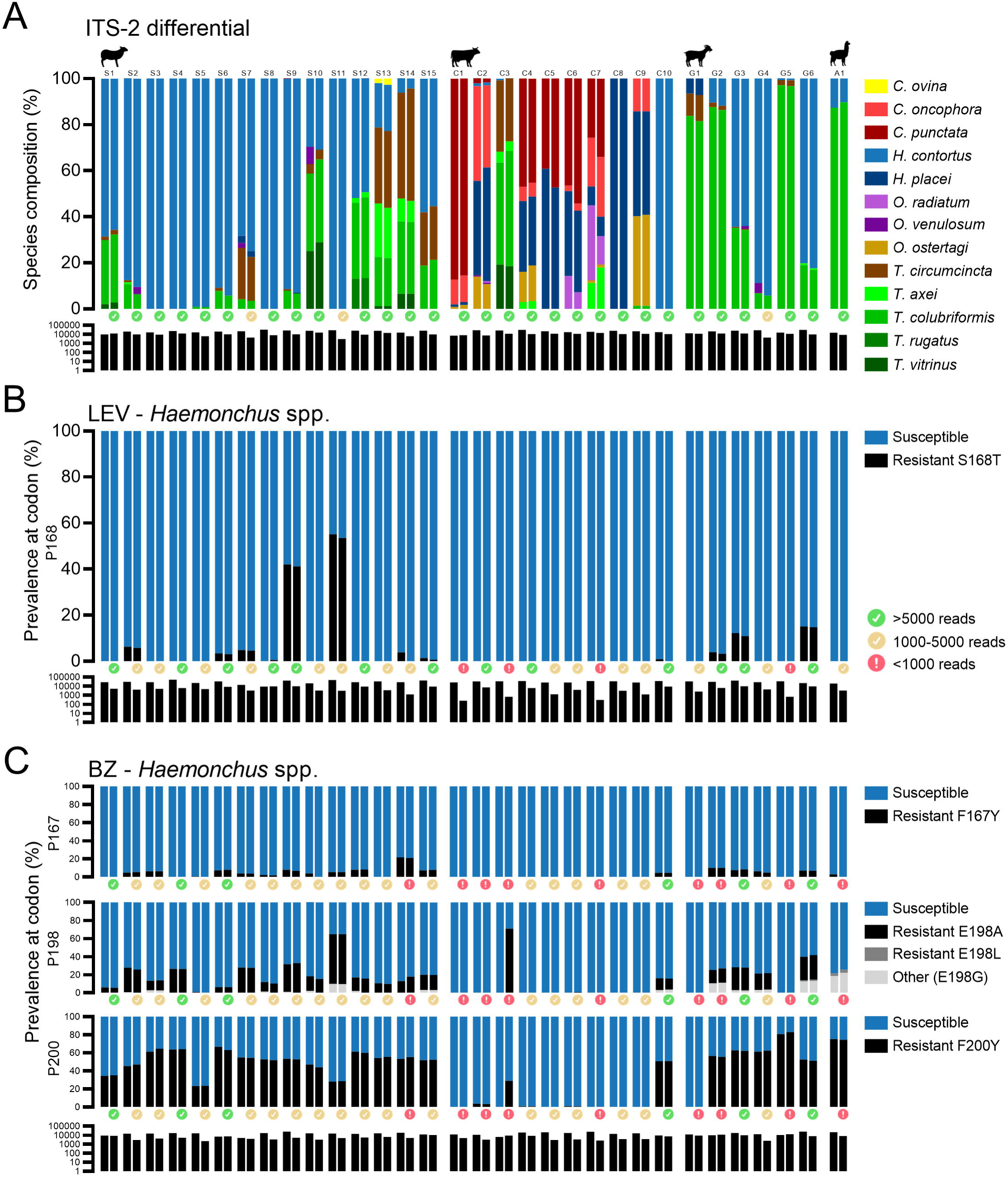
Comparison of individual and mixed amplicon metabarcoding approaches for detection and quantification of anthelminthic resistance loci and molecular larval differentiation. ITS-2 nemabiome species compositions (A), and frequencies of single nucleotide polymorphisms (SNPs) conferring levamisole (LEV) (S168T) (B) and benzimidazole (BZ) (F167Y, E198A/L/other, F200Y) (C) resistance in *Haemonchus* spp. L3. Larval culture samples (N=32) were obtained from ovine, bovine, caprine and camelid faecal samples submitted to the New South Wales Department of Primary Industries, Australia, for routine diagnostics in 2022. Each sample is presented as two bars on each bar chart based on the Illumina next generation sequencing approach used; individual amplicon metabarcoding results are displayed on the left, and results from mixing ITS-2, *acr-8* exon 4 (LEV) and *β-tubulin* isotype-1 (BZ) amplicons are displayed on the right. The total number of Illumina reads (log10) for each sample, obtained by each sequencing approach are displayed below each bar chart.

**Figure 3.**
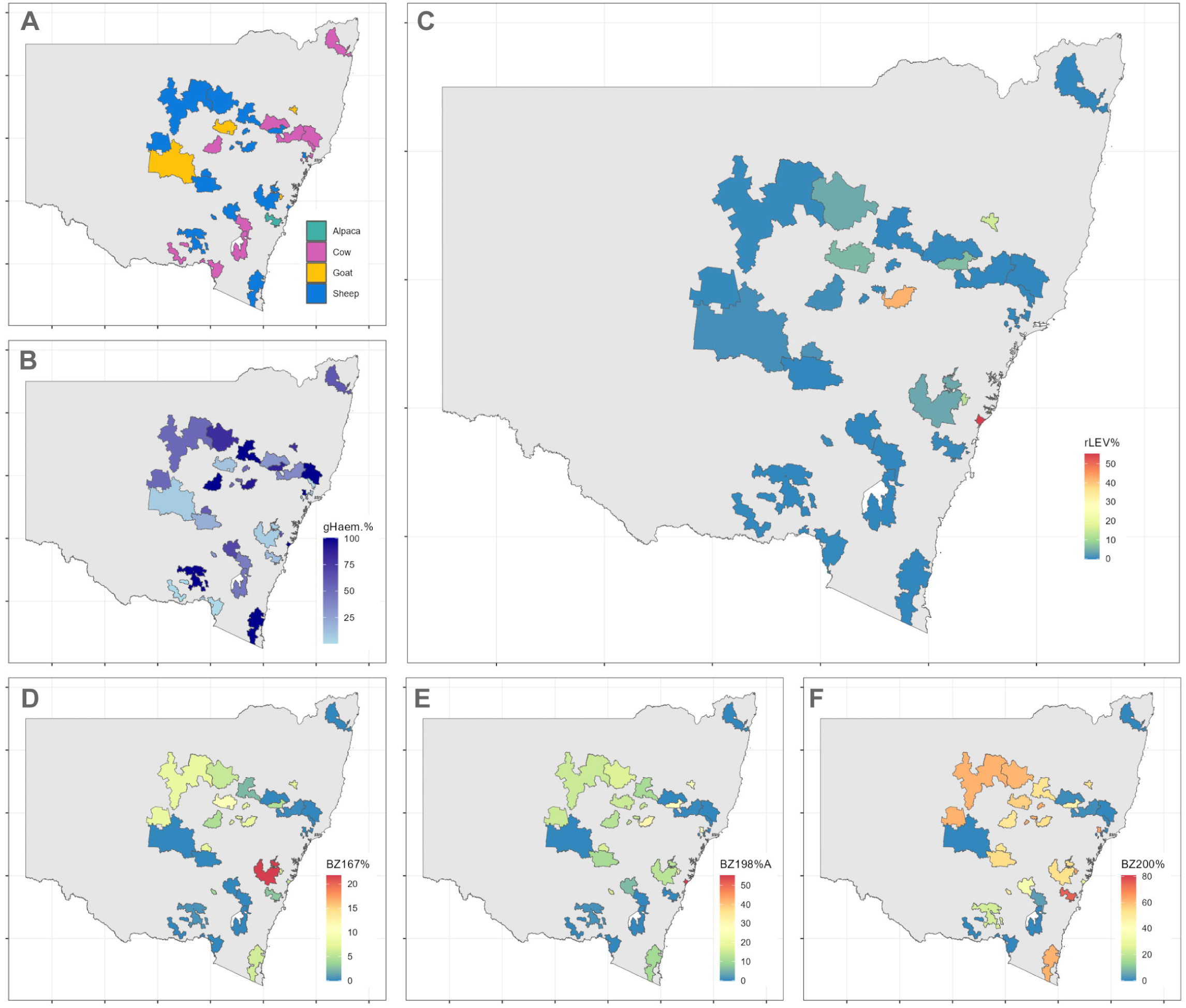
Mapping the simultaneous detection and quantification of anthelminthic resistance loci and molecular larval differentiation using metabarcoded deep amplicon sequencing. (A) Postcodes with larval culture samples (N=32) from sheep, goats, cattle and alpaca are color-coded. All samples are larval cultures and collected in 2022. (B) Detection and percentage of *Haemonchus* spp. (gHaem%) within each postcode evaluated using amplicon metabarcoded deep sequencing of ITS-2. *Haemonchus* spp. DNA was detected in all postcodes with the colour intensity proportional to the percentage from each postcode. (C) Percentage of S168T variant in *Haemonchus* spp. DNA (rLEV%) conferring levamisole resistance within each postcode evaluated using amplicon metabarcoded deep sequencing, 12/32 (37.5%) field isolates (8 sheep, 1 cattle and 3 goats). (D, E, F). Benzimidazole *β-tubulin isotype-1* metabarcoding targeting *Haemonchus* spp. F167Y (D), E198A/L (E) and F200Y (F). Shapefiles for Australia and postcodes were obtained from the Australian Bureau of Statistics (1270.0.55.001 - Australian Statistical Geography Standard (ASGS): Volume 1 - Main Structure and Greater Capital City Statistical Areas, July 2016, Australian Government).

For the ITS-2 nemabiome metabarcoding, all 32 field isolates were stage-1 SYBR-chemistry real-time PCR positive with a mean C_t_ of 17.2 (SD 4.3). Using the Illumina MiSeq (PE250) and ‘dada2’ pipeline we obtained an average sample read depth of 21,643 (SD 6,104) reads for amplicons sequenced individually, and 9,860 (SD 2,926) when mixed. A total of 68 unique ITS-2 ASVs were produced, with both individual and pooled amplicons identifying the presence of 13 trichostrongylid nematode species: *Chabertia ovina, Cooperia oncophora, Cooperia punctata, H. contortus, H. placei, Oesophagostomum radiatum, Oesophagostomum venulosum, Ostertagia ostertagi, Teladorsagia circumcincta, Trichostrongylus axei, Trichostrongylus colubriformis, Trichostrongylus rugatus* and *Trichostrongylus vitrinus* (Fig. 2A). *H. contortus* and/or *H. placei* was detected in all 32 field isolates, although contributions were minimal (<10% total reads) in some isolates (S14, C1, C3, C7, G1 and G5) (Fig. 3B). Overall, there was no statistical difference (P=0.30, two-way RM ANOVA) in species compositions between individual and pooled amplicons, even when species were in very low proportions (<2% total reads).

LEV *acr-8* exon 4 stage-1 SYBR-chemistry real-time PCRs yielded a mean C_t_ of 25.9 (SD 3.8) and were Illumina sequenced to an average sample read depth of 30,572 (SD 9,005) reads for amplicons sequenced individually, and 4,869 (SD 3,174) when mixed. A total of 56 unique *Haemonchus* spp. *acr-8* ASVs were generated with >96% identity to one of the four geographically divergent reference sequences (LS997566, CP035803, AUUS01049415, UZAF01016342). The sequence conservation of the *acr-8* exon 4 made it impossible to differentiate between *H. contortus* and *H. placei*, therefore all sequences are conservatively considered as *Haemonchus* spp. (Fig. 1D). Another 22 unique ASVs were produced and tentatively identified as either *T. circumcincta* or *Trichostrongylus* spp. based on NCBI BLAST and comparison to the reference genome data, *T. circumcincta*: Tci-*acr-8* (HQ215517) and *T. colubriformis*: Tco-*acr-8* (HQ215518) (Boulin et al., 2011) (Fig. 1D). These non-*Haemonchus* sequences were not considered further as our primers were only validated for *Haemonchus* spp. and combined they accounted for <2% of total reads (25,250/1,134,168). *Haemonchus* spp. SNP analysis revealed LEV resistant S168T in 12/32 (37.5%) field isolates (8/15 sheep, 1/10 cattle, 3/6 goats and 0/1 alpaca) with an average frequency of 13.54% (SD 18.1%) at the codon (Fig. 2B; Fig. 3C). Using the above mentioned threshold (16%), 2/32 (sheep S9 and S11) were considered unambiguously LEV resistant *Haemonchus* spp. populations. Despite the reduction in read depth, there was no statistical difference (P=0.78, two-way RM ANOVA) between the S168T resistance profiles of individual and mixed amplicons.

For BZ *β-tubulin* isotype-1 metabarcoding, stage-1 SYBR-chemistry real-time PCRs yielded a mean C_t_ of 24.7 (SD 6.5) with an average Illumina read depth of 14,859 (SD 5,108) reads for amplicons sequenced individually, and 5,881 (SD 2,888) when mixed. A total of 143 unique *β-tubulin* isotype-1 ASVs were generated with 31 assigned to the genus *Haemonchus*. SNP analysis at P167, P198 and P200 revealed similar results as observed for LEV S168T, with no statistical difference (P167: P=0.74, P198: P=0.14, P200: P=0.85, two-way RM ANOVA) observed between BZ resistant profiles for individual and mixed amplicons (Fig. 2C). The lower similarity at P198 was due to isolates with sub optimal read depth (<1,000 reads), which in all cases corresponded to having low proportions of *H. contortus* and/or *H. placei* based on ITS-2 sequencing. The F200Y variant was detected in 26/32 (81.3%) field isolates (15/15 sheep, 5/10 cattle, 5/6 goats, and 1/1 alpaca) with an average prevalence of 47.4% (SD 21.2%) when present at the codon. Detection of E198A/L was also common (total 71.8%; 15/15 sheep, 3/10 cattle, 4/6 goats, and 1/1 alpaca) with average frequencies of 20.1% (SD 16.5%) and 1.03% (SD 1.02%), respectively. Overall F167Y was also common in most hosts (total 53.1%; 11/15 sheep, 1/10 cattle, 4/6 goats, 1/1 alpaca) but had a lower prevalence at the codon (6.2%: SD 4.6%) in comparison to the other markers of BZ resistance (Fig. 3D-F). Examination of BZ resistant SNPs for other trichostrongylid nematode species identified via *β-tubulin* isotype-1 metabarcoding was not relevant to this study but can found in LabArchives (https://dx.doi.org/10.25833/9h0d-0941).

An additional 40 field L3 isolates underwent LEV *acr-8* exon 4 metabarcoding (32 ovine, 8 bovine) (Fig. 4A). Unambiguous (>16%) *Haemonchus* spp. S168T LEV resistant SNP was detected in 4/32 (12.5%) ovine isolates (S16: 18.6%, S21: 29.8%, S31: 41.8%, S45: 36.6%) with an additional 14 (43.8%) with S168T LEV resistant SNP present at frequencies of 0.8-9.9%. In total, for *Haemonchus* spp. from sheep an overall 6/47 (12%; 95%CI = 5.6% to 25.6%) were considered S168T LEV resistant ovine populations across NSW (Fig. 4B). Only a single bovine (1/8) had *Haemonchus* spp. S168T LEV resistant SNP (1.2%); overall no *Haemonchus* spp. population from NSW cattle (0/18) had the S168T LEV resistant SNP at frequency >16%.

**Figure 4.**
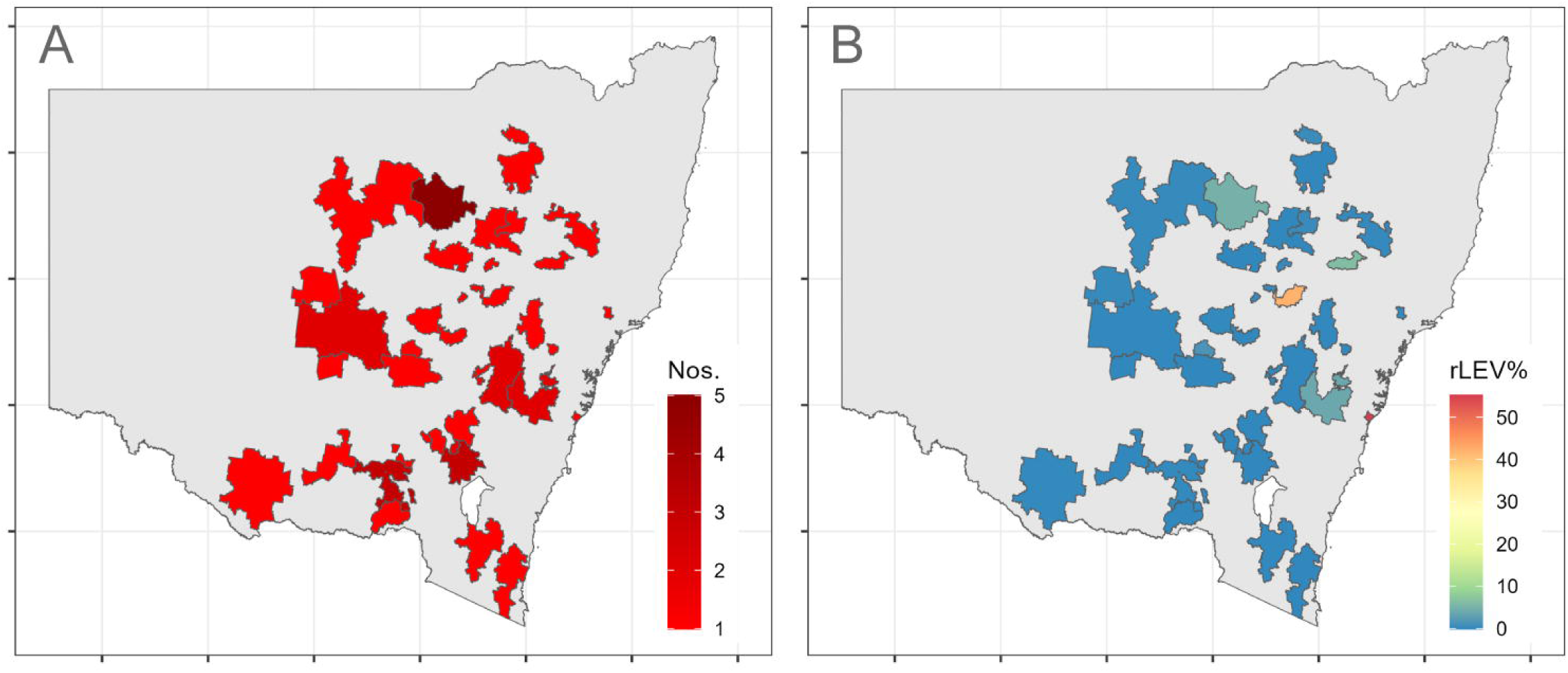
Distribution of *Haemonchus contortus* S168T SNP conferring lowered efficacy to levamisole across New South Wales, Australia. (A) Postcodes with sheep samples (N=47) are highlighted red and the colour intensity is proportional to the number (1 to 5) of samples from each postcode. All samples are larval cultures and collected in 2022. (B) Percentage of S168T variant in *H. contortus* DNA (rLEV%) within each postcode evaluated using amplicon metabarcoded deep sequencing. Shapefiles for Australia and postcodes were obtained from the Australian Bureau of Statistics (1270.0.55.001 - Australian Statistical Geography Standard (ASGS): Volume 1 - Main Structure and Greater Capital City Statistical Areas, July 2016, Australian Government).

## 4. Discussion

We describe a powerful new mixed amplicon metabarcoding and sequencing approach for identifying parasitic nematode populations and screening polymorphisms associated with *Haemonchus* spp. resistance to two major classes of anthelmintics. To our knowledge, this constitutes the first multi-drug and multi-species molecular diagnostic developed for helminths of veterinary importance. Until now, deep amplicon next generation sequencing of anthelmintic resistance polymorphisms had only been applied to the *β-tubulin* isotype-1 gene for detection of SNPs at codons 167, 198 and 200, conferring BZ resistance in trichostrongylid nematodes of ruminants (Avramenko et al., 2019; Sargison et al., 2019; Avramenko et al., 2020; Melville et al., 2020; Evans et al., 2021). The recent discovery of a non-synonymous variant, S168T, in *H. contortus acr-8* exon 4, demonstrated to be strongly correlated with LEV resistance across multiple datasets (Doyle et al., 2022), prompted us to design and validate a LEV *acr-8* metabarcoding technique. Detection of the S168T SNP previously utilised allele-specific (AS) and droplet-digital (dd) PCR, which is lower-throughput than metabarcoding and the AS-PCR performed suboptimally for Australian isolates, most likely due to genetic variation in the primer binding site (Antonopoulos et al., 2022; Baltrušis et al., 2023). Our LEV *acr-8* metabarcoding technique successfully detected proportions of S168T in Australian LEV resistant *H. contortus* GWBII and enables state or country wide surveillance.

In the face of growing multi-drug resistance, molecular tools like LEV *acr-8* and BZ *β-tubulin* isotype-1 metabarcoding would be most useful if incorporated into a single assay for detection of SNPs associated with anthelmintic resistance across multiple drug classes. Consequently, we validated an approach in which ITS-2, LEV *acr-8* exon 4 and *β-tubulin* isotype-1 amplicons were mixed into a single sequencing sample for metabarcoding, resulting in successful detection of resistant SNP allele proportions and GI nemabiomes to the level of standard individual metabarcoding techniques. Our approach is not only cost effective (in our case for 384-indexed 250PE Illumina run: ∼$20AUD, ∼13USD or ∼12EUR for all three targets per isolate), but it is highly scalable due to the utilisation of robotics and increased availability of high-throughput sequencing technologies including portable sequencing devices (Liou et al., 2020; Whitford et al., 2022). It would overcome many of the limitations of current *in vivo* and *in vitro* phenotypic resistance tests, e.g. long turn-around time for results (Kotze et al., 2020) and insensitivities to low levels of resistance (<25%) (Martin et al., 1989). Most importantly, implementation in the field would be beneficial in providing information as to what alternative drugs remain effective on a property, informing management decisions and promoting the longevity of available anthelmintics: so far representing the most ‘ideal’ molecular diagnostic tool currently described, based on the key criteria outlined by the Consortium for Anthelmintic Resistance and Susceptibility (CARS) (Kotze et al., 2020).

Despite the benefits of a mixed amplicon metabarcoding and sequencing tool, interpreting the functional connection between SNP frequencies and the actual phenotypic status of GIN L3 populations currently presents a challenge. An arbitrary 10% threshold was considered for BZ 167, 198 and 200 SNPs by Barrère et al. (2013a) for farms with *H. contortus*; backed by their previous study using FECRTs with −26% to 54% BZ efficacy on those farms (Barrère et al., 2013b). Our LEV threshold (16%) is based on WGS of resistant *H. contortus* GWBII isolate with 41% LEV efficacy based on total worm count (Love et al., 2003; Kotze et al., 2018; Ruffell et al., 2018). In the case of LEV *acr-8* S168T it is possible that the marker is unlikely to explain all cases of phenotypic resistance in *H. contortus* (Baltrušis et al., 2023). A second QTL in *H. contortus* MHco3/18 cross following LEV selection implicates the *lev-1* gene region on chromosome IV as well (Doyle et al., 2022). The potential mechanistic role of S168T in LEV resistance is an aspect that should be explored in future work, which could make use of the *Xenopus* oocyte system instrumental in deciphering AChR subunit composition (Boulin et al., 2011). Nonetheless, our detection of the S168T SNP in geographically separate and genetically divergent populations to those described in the initial studies (Doyle et al., 2022; Antonopoulos et al., 2022) further validates the efficacy of S168T as a marker of LEV resistance.

The GWBII isolate has previously been shown to behave as two distinct subpopulations in larval development assays (LDA): a low-level LEV resistance fraction (∼45%) that shows approximately 2-fold resistance to LEV at the IC50, and a highly resistant fraction (∼55%) that is unaffected by the drug at concentrations over 100-fold greater than the IC50 of susceptible isolates (Ruffell et al., 2018). The lower proportion of the S168T allele detected via WGS, in comparison to the proportion of GWBII highly resistant via LDA, could potentially be explained by the heterozygosity of individual L3, as explored in BZ SNPs (Barrère et al., 2012; Kotze et al., 2012). This is of interest to our understanding of LEV resistance as the MHco18(UGA2004) isolate also displayed three distinct sub-populations in relation to S168T when examined by AS-PCR: a large minority, ∼20%, were homozygous for S168T, in addition to a large minority ∼20% were homozygous for S168, with the remainder being heterozygous at codon 168 (Antonopoulos et al., 2022). These findings, in addition to our own described here, indicate that LEV resistance could exist on a spectrum, with high level resistance conferred by homozygous S168T, whereas low to moderate resistance may be conferred by heterozygosity at this position. Further work is necessary to clarify this, however, in the intermediate term, this suggests that a percentage proportion of S168T within a population could be used to effectively track the emergence of resistance within a farm.

Of the NSW field isolates we tested, S168T was detected in the majority of *Haemonchus* spp. populations from sheep and goats, but rarely at a frequency greater than 16%. Overall, BZ resistance was far more prevalent than LEV resistance in *Haemonchus* spp., particularly at codons 167 and 200, confirming that LEV is still an important anthelmintic class for small ruminants in NSW (Playford et al., 2014), but requires more regular wide-scale surveillance for sustainable continued use in the field. Our results confirmed that in *Haemonchus* spp., LEV and BZ resistance is relatively uncommon in NSW cattle, however caution needs to be exercised in particular with the introduction of new combination pour-on drenches (Kotze et al., 2020). Finally, we show that S168T is present in geographically separate and genetically divergent populations of *Haemonchus* spp.

## 5. Conclusions

In order to interpret *H. contortus* LEV *acr-8* metabarcoding of field isolates in the absence of phenotypic data, we propose a working hypothesis to be considered in future field studies. Presence of 16% S168T in *H. contortus* populations reflects LEV efficacy <70%, in which LEV should no longer be used. Detection of the S168T allele at a frequency below this threshold indicates emerging LEV resistance that may not yet be phenotypically detectable, but that would be expected to increase rapidly to clinical resistance with ongoing selection. The findings described herein, and the multi-drug and multi-species molecular diagnostic approach used, is significant as it opens the way towards the use of large-scale sequencing as a surveillance technology, the results of which can be translated into concrete recommendations for the livestock sector.

## Acknowledgements

We thank Brendan Sharpe (Invetus, Armidale, Australia) for donating the *Haemonchus* spp. isolates (Kirby, GWBII and Grieve) as cryopreserved larvae and Andrew Kotze (CSIRO) for discussion and insight about levamisole resistance in nematodes. This work was in part funded by the grant from the McGarvie Smith Institute, Australia. The authors would like to thank Denis and Erika Pidcock for research funds and providing EKF’s PhD stipend. EKF’s travel to the University of Glasgow was funded by the Grants-in-Aid and Postgraduate Research Support Scheme, University of Sydney. We acknowledge the Sydney Informatics Hub (University of Sydney, Australia) for ongoing technical support and access to the high-performance computing facility, Artemis.

