## Supplementary material for "A mixed amplicon metabarcoding and sequencing approach for surveillance of drug resistance to levamisole and benzimidazole in *Haemonchus* spp": Table 1

Table 1. | **Amplification primers for the 168 SNP in *acr-8* exon 4 of *Haemonchus* spp. adapted for Illumina indexing and metabarcoding**

| Name | Primer ID | *Sequence 5’-3’ |
| --- | --- | --- |
| LEV1 | S1127 | *TCG TCG GCA GCG TCA GAT GTG TAT AAG AGA CAG* GGT AAC TGC CGC ACA TCT AAG AG |
| LEV2 | S1128 | *GTC TCG TGG GCT CGG AGA TGT GTA TAA GAG ACA G* GCC GAT GGT GAG CCT CAT ATT AC |
| LEV3 | S1135 | *TCG TCG GCA GCG TCA GAT GTG TAT AAG AGA CAG* GTG ATT TCG TGC AGA GAT AGG |
| LEV4 | S1136 | *TCG TCG GCA GCG TCA GAT GTG TAT AAG AGA CAG* GTG ATT TCC TGT AGA GAT AGG |
| LEV5 | S1137 | *TCG TCG GCA GCG TCA GAT GTG TAT AAG AGA CAG* GTG ATT TCS TGY AGA GAT AGG |
| LEV6 | S1138 | *TCG TCG GCA GCG TCA GAT GTG TAT AAG AGA CAG* GGW ARC TGY CGC ACA TCT AAG AG |
| LEV7 | S1139 | *TCG TCG GCA GCG TCA GAT GTG TAT AAG AGA CAG* GGA AGC TGC CGC ACA TCT AAG AG |
| LEV8 | S1140 | *TCG TCG GCA GCG TCA GAT GTG TAT AAG AGA CAG* GGT AGC TGT CGC ACA TCT AAG AG |

Note: *sequence in Italic indicates the Illumina overhang adapter sequences (LEV1, LEV3-LEV8 are used in combination with LEV2)
